# CABO-16S – A Combined Archaea, Bacteria, Organelle 16S database for amplicon analysis of prokaryotes and eukaryotes in environmental samples

**DOI:** 10.1101/2024.10.23.619938

**Authors:** Eryn M. Eitel, Daniel Utter, Stephanie Connon, Victoria J. Orphan, Ranjani Murali

## Abstract

Identification of both prokaryotic and eukaryotic microorganisms in environmental samples is currently challenged by either the burden of additional sequencing required to obtain both 16S and 18S rRNA sequences or the introduction of multiple biases induced by the use of “universal” primers. Organellar 16S rRNA sequences are automatically amplified and sequenced along with prokaryote 16S rRNA, and may provide an alternative method to identify eukaryotic microorganisms. CABO-16S combines bacterial and archaeal sequences from the SILVA database with 16S rRNA sequences of plastids and other organelles from the PR2 database to enable identification of all 16S rRNA sequences. Comparison of CABO-16S with SILVA 138.2 results in equivalent taxonomic classification of mock communities and increased classification of diverse environmental samples. In particular, identification of phototrophic eukaryotes in shallow seagrass environments, marine waters, and lake waters was increased. CABO-16S also provides the framework to add curated datasets of specialized sequences for further classification of clades which are not currently included in other databases. Addition of sequences obtained from Sanger sequencing of methane seep sediments and curated sequences of the polyphyletic SEEP-SRB1 clade resulted in differentiation of syntrophic and non-syntrophic SEEP-SRB1 in hydrothermal vent sediments. Such additions may simplify analysis of communities contributing to the anaerobic oxidation of methane, and highlight the potential benefit of amending existing training sets with curated sequences when studying extreme or unique environments underrepresented in existing databases.

## Introduction

Life on earth is composed of bacteria, archaea, and eukaryotes. All three branches of Domains significantly influence ecosystem function [1] and while the impact and response of humans and other macro-organisms to their environment can often be directly observed [2-4], the role of unseen microorganisms is equally important [5-7]. Microorganisms generally refer to any organism not individually visible to the human eye and are usually smaller than 50 μm. This includes bacteria, archaea, viruses, fungi, and protists, with protists referring to unicellular eukaryotes which can be free-living or form colonies, but do not have differentiated cellular functions. Over the last 20 years, high-throughput sequencing of small subunit (SSU) ribosomal RNA (rRNA) has enabled study of the microbial ecology in terrestrial [8] and marine [9] environments and facilitated understanding of the microbiome in plants [10] and animals [11]. Identification of prokaryotes was pioneered using the 16S rRNA gene [12] while the homologous 18S rRNA gene was optimized for eukaryote identification [13] and the internal transcribed spacer (ITS) region proved best for fungi [14].

Amplification of both prokaryotic and eukaryotic microorganisms with a single PCR reaction would be ideal and could reduce amplicon library preparation costs two to three-fold compared to the use of separate primers for 16S and 18S rRNA gene sequencing. Although some “universal” primers (515f/926r) can amplify eukaryotic 18S rRNA along with 16S rRNA [15], simultaneous and accurate analysis of both eukaryotes and prokaryotes from 16S and 18S rRNA is challenging. First, mismatches between the primers and its template may be more common when attempting to amplify a broader target group, and indeed, in mock communities only a single mismatch with the reverse primer resulted in a 3-8 fold underestimation of eukaryotes [16]. Second, 18S sequences are typically 160-180 bp longer than 16S sequences and both PCR and sequencing is biased against longer amplicons [17]. Natural samples may contain long 18S sequences [18] or higher proportion of dinoflagellates which tend to have mismatches suggesting that underestimation of eukaryotes with 515f/926r primers may be significantly worse in some natural samples [16]. Finally, 16S rRNA gene copy numbers range from 1 to 15 in most bacteria and average only one copy in a majority of archaeal phyla [19, 20] while 18S gene copy numbers can vary between 1 and 12,000 in phytoplankton [21]. Although 18S gene counts may be significantly correlated with biovolumes, they cannot be reliably used to determine relative taxonomic abundances [22].

Plastids, eukaryotic organelles originating from ancestral cyanobacterial endosymbionts, contain 16S rRNA copies which are taxonomically linked to their host organism rather than their cyanobacterial past. Plastid 16S rRNA gene copies are more constrained than 18S rRNA gene copies. The number of plastids, such as chloroplasts, vary less with cell size than 18S rRNA. Although environmental factors can still influence chloroplast counts, particularly in large taxa [23-25], within the chloroplasts there are typically 1-2 copies of the 16S rRNA gene. Together, this results in similar ranges of plastidal 16S rRNA gene counts and bacterial or archaeal 16S rRNA gene counts. Furthermore, phytoplankton taxonomy determined with 18S rRNA gene sequences is comparable to 16S rRNA [25, 26] and in some cases plastid 16S rRNA may provide better phylogenetic resolution than 18S rRNA [27, 28].

Plastidal 16S rRNA analysis appears to provide a pathway for identification of bacterial, archaeal, and most photosynthetic eukaryotic microorganisms relevant in marine and terrestrial environments without the use of multiple primers or having to account for the non-uniform biases introduced during attempts to quantify both 16S and 18S rRNA with “universal” primers. As a part of routine 16S rRNA analysis, plastidal 16S are automatically amplified and sequenced along with prokaryote 16S rRNA. However, identification of plastid sequences is not currently possible as a curated reference database that includes bacterial, archaeal, and plastidal sequences for taxonomic identification is lacking. Tedious manual verification is required to demonstrate that unidentified sequences are in fact plastids. If not annotated, plastid sequences can complicate sample analysis by artificially inflating the fraction of unidentified taxa in a sample, potentially leading to erroneous inference of ‘novel’ taxa. The SILVA database (v138.2) [29] contains a comprehensive record of 16S rRNA gene sequences and is continuously updated to include recently identified taxa from novel environments, however only broad identification of chloroplasts is possible with no ability to provide further taxonomic identification. On the other hand, the database PR2 [30] may contain the most comprehensive plastidal 16S rRNA record, but with a focus on protists, limited bacterial and archaeal sequences are supplied. With the CABO-16S (Combined Archaeal, Bacterial, and Organelle for 16S) database, we combine the bacterial and archaeal sequences from SILVA with the 16S sequences of plastids and other organelles from PR2 to enable efficient identification of all 16S sequences. A framework is provided for CABO to be updated with future releases of SILVA and PR2 or users can include sequences particular to their own area of interest. Taxonomic classification is compared between CABO-16S and SILVA-132.1 in mock microbial communities and published datasets from mammalian guts, deep-sea seep sites, seagrass beds, alkaline lakes with high abundances of microalgae, and terrestrial soils. Finally, addition of custom sequences to CABO-16S can allow identification of specialized taxonomic groups, even increasing our understanding of polyphyletic groups which are difficult to constrain within the current SILVA structure. This may allow deeper understanding of complex environments, such as deep-sea hydrothermal vents.

## Methods

### Aggregation of reference databases

To build the CABO-16S database, sequences from the most recent version of Silva available (138.2) were downloaded (SILVA_138.2_SSURef_NR99_tax_silva.fasta.gz) along with mapped taxonomy (taxmap_slv_ssu_ref_nr_138.2.txt.gz) and quality values (SILVA_138.2_SSURef_Nr99.quality.gz). All sequences with a pintail value < 50 or an alignment quality value < 75 were removed. Sequences identified as Chloroplast, Mitochondria, and Eukaryotes were removed, except for 100 randomly selected Eukaryotes which were added back in as an outgroup. The taxonomy of the Eukaryotic outgroup was only retained at the Phylum level. The prokaryotic taxonomy was cleaned up, particularly at the Species level to remove naming schemes based on organism host, sample collection, unclear bacterium groupings, or repetition of genus (ie “Genus sp.”). For direct comparison, a simplified SILVA database was constructed using identical methods with the exception of retaining sequences identified as Chloroplasts.

Plastids, apicoplast, mitochondrion, and chromatophores sequences were added to the CABO-16S database from the PR2 database (v 5.0.0) with acquistion using the R package ‘pr2database’ (https://pr2database.github.io/pr2database/articles/pr2database.html). To match the 7 rank orders found in SILVA taxonomy, the ranks of Supergroup and Subdivision were dropped from PR2 sequences. Finally, custom 16S rRNA sequences obtained from Sanger sequencing of methane seeps (https://doi.org/10.6084/m9.figshare.27288090) were combined with the selections from SILVA and PR2 to form the basis of the CABO-16S dataset.

The CABO-16S and simplified SILVA 138.2 training sets were made using recommendations from DECIPHER [31] and based on the IDTAXA algorithm [32]. Briefly, oversampled groups were randomly subset to 100 sequences before training over three iterations with the LearnTaxa function. Kmer length was set to 8nt to match RDP and QIIME2 defaults. Note that full-length 16S rRNA reference sequences were used for training; truncation to the amplicon window may slightly improve accuracy (e.g. [33]) but at the cost of potentially generating ambiguity [34]. Thus, we present full-length sequences and perform comparisons from full-length sequences and leave the choice of truncation to users.

### Classification of benchmarking datasets

Taxonomic classifications using both CABO-16S and SILVA-132.1 were compared in published 16S rRNA sequences from a broad range of sources including both mock communities of known bacterial isolates and environmental samples. For all compared samples, the V4-V5 region of the 16S rRNA gene was amplified using archaeal/bacterial primers (515f/926r) and sequenced on the Illumina MiSeq platform. In environmental datasets with more than 5 samples, an arbitrary set of subsamples was selected for taxonomic classification comparison. Published metadata and accession numbers or data sources are combined in Table S1; full methodological details regarding the generation of the datasets can be found in the original papers.

All downloaded raw sequences were processed identically, except for Needham and Fuhrman (2016) data for which the already-analyzed OTU sequences and observation matrices were downloaded and used directly. A reproducible workflow (https://github.com/emelissa3/CABO-16S commit 472d7fc) reports the full details and parameters used for generating amplicon sequence variants (ASVs) from the raw FASTQ files available on NCBI SRA. Briefly, primers were removed using Cutadapt [35] then sequences were trimmed (240f/200r), merged with a 12 bp overlap, denoised, and aligned using DADA2 [36]. Chimeras were removed and taxonomy was assigned by the IdTaxa function from IDTAXA [32]. IdTaxa parameters were default except for using a confidence threshold of 40%, as this produced classifications most directly comparable to the default QIIME2 classifications based on unpublished comparisons. ASV counts and taxonomic assignments are reported in Table S2, and ASV sequences are in Data_ASVs.fa.

## Results/Discussion

CABO-16S is a practical unification of commonly used 16S rRNA databases to provide a single database that can be easily expanded by users to incorporate database updates or incorporate unpublished sequences. The 389,144 bacterial and 19,213 archaeal 16S rRNA sequences from SILVA 138.2 [29] were used as the initial backbone of CABO-16S database, along with a random 100 Eukaryota sequences from SILVA retained as an outgroup. These were combined with 8,540 16S rRNA sequences from organellar 16S rRNA genes from the PR2 database [30]. Finally, custom sequences can be combined to maximize resolution into target groups; here we added a set of unpublished full-length 16S rRNA sequences obtained from Sanger sequencing of methane seep sediments and a curated list of representative SEEP-SRB1 sequences [37].

### CABO-16S annotates previously unclassified ASVs with comparable accuracy

CABO-16S was compared against SILVA 138.2 using a combination of previously published datasets representing diverse systems, including both mock communities and environmental samples (Table S1). The dataset consists of well-characterized benchmarking sets based on mock communities, mammalian guts, and residential soils [38, 39], boreal forest soils [40], seagrass roots, leaves, and surrounding sediment [41], deep-sea sediments from cold methane seeps [42], hydrothermal vent sediment [43], marine water with abundant phytoplankton communities [25], and finally, water from a closed basin lake dominated by the microalgae Picocystis [44]. The combined dataset consists of 64,402 ASVs with individual datasets ranging from 45-32,090 ASVs.

The CABO-16S database outperformed the unamended SILVA 138.2 database in terms of the total number of ASVs receiving taxonomic assignment across all taxonomic levels (Figure 2). The largest differences were the datasets with the most phototrophic eukaryotes, such as the shallow seagrass environments, marine waters, and lake waters. For example, in the marine water dataset, CABO-16S allowed classification of approximately 10% more ASVs than SILVA 138.2 at the phylum level (Figure 2c). Other datasets differed little, suggesting that the inclusion of the PR2 database’s organelle 16S sequences did not meaningfully impact the ability of the classifier to continue to accurately predict bacterial and archaeal taxonomy. The only notable exception was the marine water dataset had slightly higher classification rates in SILVA 138.2 than with CABO-16S, which we attribute to the increased number of phytoplankton orders from PR2 vs. the singular “Chloroplast” label in SILVA. The vast majority of ASVs could not be classified at the species level in either dataset, although this may be partly due to relatively few reference sequences being labeled down to the species level, particularly for microbes endemic to non-human environments.

**Figure 1.**
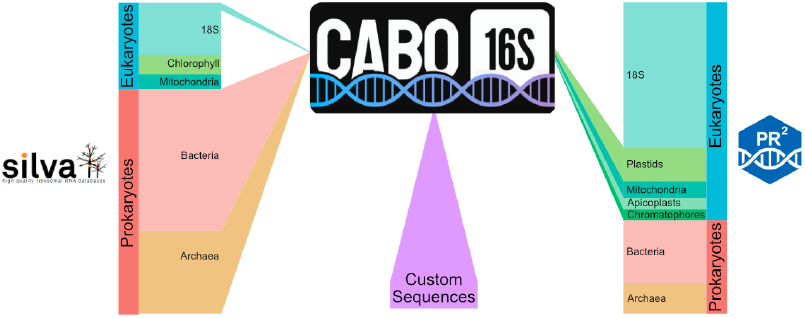
Schematic showing the combination of SILVA, PR2, and custom sequences to form the CABO-16S database.

**Figure 2.**
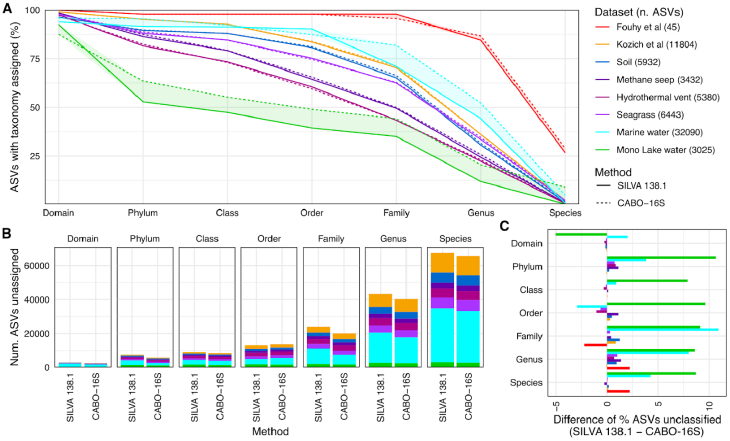
The CABO-16S database improves taxonomic classification in environmental datasets with phototrophic eukaryotes. A) Percentage of ASVs classified at a given taxonomic level (x-axis) by dataset (colored lines) for SILVA 138.2 (dashed) and CABO-16S (solid). The lines continuously decrease as ASVs lacking annotation at higher ranks, e.g., Domain, are by definition also lacking annotation for lower ranks, e.g., Species. B) Absolute number of ASVs lacking classification, as in A. C) The difference in percentage unclassified between the two databases. Positive percentages reflect CABO-16S annotating more ASVs than SILVA 138.2, vice versa for negative percentages.

Both CABO-16S and SILVA classifications reveal a distinction between two types of ambiguity which can prevent taxonomic annotation. Ambiguity is the most commonly considered form of precision, where a sequence is placed intermediate between two or more reference taxa and thus cannot be assigned to a single taxon at a chosen confidence threshold (40% in this study). IDTAXA and other similar classifiers handle such occurrences by classifying the sequence to the lowest common level of the competing reference taxa, and sometimes adding a prefix of ‘unclassified’ to the conflicted taxonomic rank. Conversely, a sequence may be confidently assigned to a single taxon, however taxonomy may still be lacking at a given rank if the reference sequence lacks annotation at that rank. Such a scenario affects many uncultured lineages, e.g. the Candidate Phylum Radiation family SR1 (phylum Patescibacteria) has no genus or species assignments in SILVA 138.2, all 121 sequences are annotated only to the family level. Thus, a SR1 ASV lacking a genus classification is not due to classifier uncertainty but rather taxonomic uncertainty. Furthermore, some lineages can include both sources of uncertainty, e.g., in SILVA 138.2 the family Desulfosarcinacae has 53 sequences labeled to the species level, 676 labeled to the genus level, and 345 labeled to the family level. Therefore, Desulfobacteraceae ASVs lacking genus-level annotation could be due to a close similarity to a set of sequences labeled only to the family level (taxonomic ambiguity) or by being indistinguishable from different genera (classifier ambiguity). Thus, we distinguish between the two using IDTAXA’s convention of prepending ‘unclassified_’ to situations of classifier ambiguity and an additional convention of prepending ‘unspecified_’ to the lowest taxon level for situations of reference sequence ambiguity.

Of the classified sequences, CABO-16S and SILVA produced similar community compositions in most datasets (Figure 3). Indeed, as the non-cyanobacterial portion of CABO-16S archaea and bacterial sequences are exactly shared with SILVA 138.2, this agreement is expected. However, in datasets with phototrophic eukaryotes (e.g. seagrass, marine and lake water column datasets), the CABO-16S database allowed classification of eukaryotic chloroplasts which accounted for nearly 50% of the reads in some samples (Mono Lake dataset, Figure 3). The marine water dataset had by far the greatest phytoplankton diversity of any of the example sets (Needham and Fuhrman, 2016); much of this diversity could be assigned a taxonomic label with CABO-16S. In the Mono Lake dataset, the remaining unclassified diversity could be attributed to phytoplankton mitochondrial sequences by manual blasting vs. NCBI. While the current PR2 database includes approximately 1842 mitochondrial sequences, the vast majority (1,782 or 96.7%) belong to Opisthokonta, with only 22 sequences spanning Archaeplastida (plants and many algae). Although mitochondria are not found in all eukaryotic cells [45], we anticipate the future expansion of PR2 to include more mitochondrial 16S from plant and algal lineages will ameliorate this problem.

**Figure 3.**
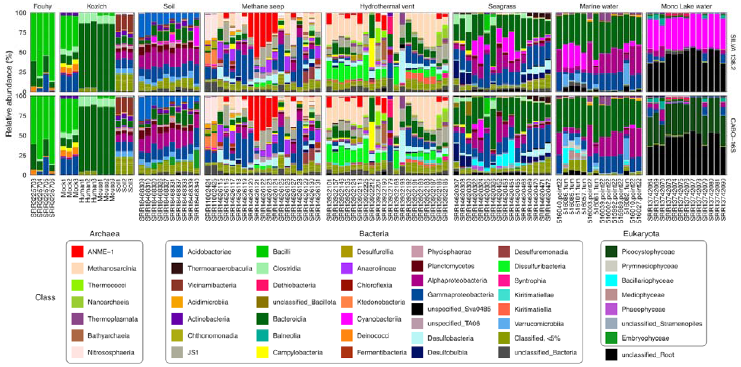
Composition of each dataset with CABO-16S vs. original SILVA 138.2. ASVs were aggregated to the class level (colors). Classes with at least 5% in any sample are shown. ASVs that could not be assigned a specific class were similarly aggregated at the lowest annotated rank. Remaining ASVs with <5% relative abundance were grouped into a single category.

### Enhanced identification of phototrophic eukaryotes

We tracked ASV classification through each dataset for phototrophic eukaryotes to elucidate points of difference between the databases (Figure 4). For the coastal marine water dataset [25] undergoing a eukaryotic phytoplankton bloom, SILVA 138.2 was able to classify the bacterial community accurately, but a large fraction of the reads, presumably eukaryotic, were not assigned at the Domain level or simply annotated as Chloroplast at the Family level’ However, with the CABO-16S dataset these same plastid ASV sequences received further taxonomic assignment (Figure 4). Notably, the diversity of sequences did not always allow for unambiguous taxonomic assignment to lower levels, i.e. tracing the taxonomic assignment in Figure 4 reveals many ambiguities arise at the class or order level. Some taxon ranks include ‘_X’ suffixes, which are placeholder intermediate used by PR2 akin to *‘Incertae Sedis’* used in other taxonomies. What taxonomic assignment can be obtained through plastid classification is useful, however, as the major phytoplankton groups (e.g. diatoms, dinoflagellates, cryptophytes, etc) are distinguished.

**Figure 4.**
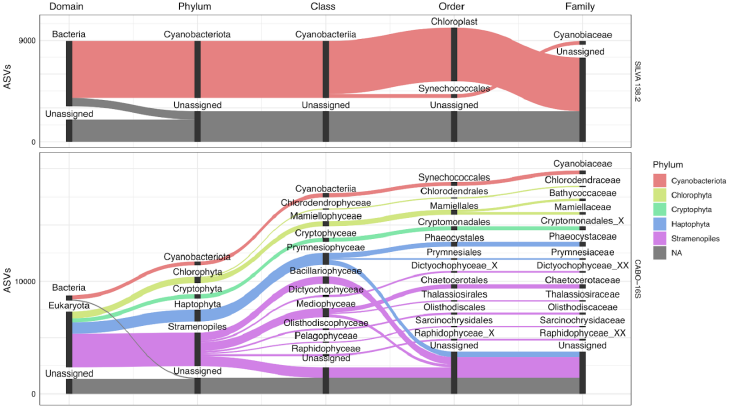
CABO-16S resolves both eukaryotic and bacterial phytoplankton in the coastal marine dataset. Alluvial plots for SILVA 138.2 (top panel) and CABO-16S (bottom panel) tracking ASV classification across taxonomic ranks from Domain (leftmost) to Family (rightmost) for families with at least 100 ASVs. For each rank, bars represent the different taxa classified with the size of each bar scaled to reflecting the number of ASVs. Whitespace between bars is for ease of visualization. Flows connecting ranks are colored based on the phylum-level classification of those ASVs. Only ASVs related to cyanobacteria or plastid sequences based on either database are shown.

### Custom sequences increase identification of polyphyletic clades

Addition of custom sequences to CABO-16S can increase taxonomic classification of species not currently included in either the SILVA or PR2 databases. We added sequences obtained from Sanger sequencing of methane seep sediments and a curated list of representative SEEP-SRB1 sequences [37]. SEEP-SRB1 is a polyphyletic clade of sulfate-reducing bacteria [46, 47], including known syntrophic partners of ANME during the anaerobic oxidation of methane (AOM), such as SEEP-SRB1a and SEEP-SRB1g, and other non-syntrophic members (SEEP-SRB1b, SEEP-SRB1c, SEEP-SRB1d, SEEP-SRB1e, and SEEP-SRB1f). Although currently identified as a genus level clade in SILVA 138.2, this is be an overly simplified grouping of these organisms. Indeed, while some members, such as SEEP-SRB1g and SEEP-SRB1c, have been described as species level clades, others such as SEEP-SRB1a, are more accurately described as genus-level clades. Further complicating SEEP-SRB taxonomy is the asymmetric phylogenetic distance between SEEP-SRB1 subgroups - for example, SEEP-SRB1g and SEEP-SRB1a may reside in different orders based on genomic trees [37, 48-50]. While rectifying phylogenetic distance with taxonomic classification is beyond the scope of this work, we note that such conflicts between historical naming conventions are unfortunately common in environmental microbiology and difficult to resolve. However, expanding databases with precisely named groups offers a means to circumvent these discrepancies. Thus, we added the representative sequences for SEEP-SRB subgroups as a “species” of SEEP-SRB1, except for SRB1g which was added as a “species” of *Desulfosudis*, the taxonomic designation from SILVA 138.2 for sequences most similar to SRB1g.

The inclusion of these additional SEEP-SRB sequences in the CABO-16S database resolved a portion of the environmental SRB1 group ASVs to their respective subgroups (Figure 5). In the methane seep and hydrothermal vent datasets, the improved resolution revealed varying distributions of the different SEEP-SRB1 subgroups (Figure 5A). In the vent dataset, only a subset of samples harbored the syntrophic Seep-SRB1a vs. the non-syntrophic Seep-SRB1d, a distinction that was not resolvable with the default SILVA 138.2 database. Further differences in taxonomic assignment become clear when tracking how ASVs annotated in SILVA as SEEP-SRB1, unclassified_Desulfosarcinaceae, or *Desulfosudis* are classified by CABO-16S compared to SILVA 138.2 (Figure 5B). While a relatively small proportion of the total ASVs from the entire dataset were differently classified by CABO-16S (Figure 5B), based on Figure 5A, the differences in classification are significant in particular environments such as the sedimented hydrothermal vents included in this study. Interestingly, a few ASVs classified as Desulfosarcinaceae with SILVA 138.2 were unclassified at higher ranks with CABO-16S (e.g., unclassified_Desulfobacterales) or differently classified (Figure 5B). We attribute such variance due to inter-run variance in the IDTAXA algorithm, as it randomly subsamples kmers for each run and so a small portion of ASVs at the edge of classification confidence threshold receive different classification between runs [32]. In support of this observation, flipping the analysis to include ASVs classified as SEEP-SRB1, Desulfosarcinaceae, or LCP-80 with CABO-16S produced a similar result of overall agreement with some ASVs classified with CABO-16S as Desulfosarcinaceae being unclassified with SILVA 138.2.

**Figure 5.**
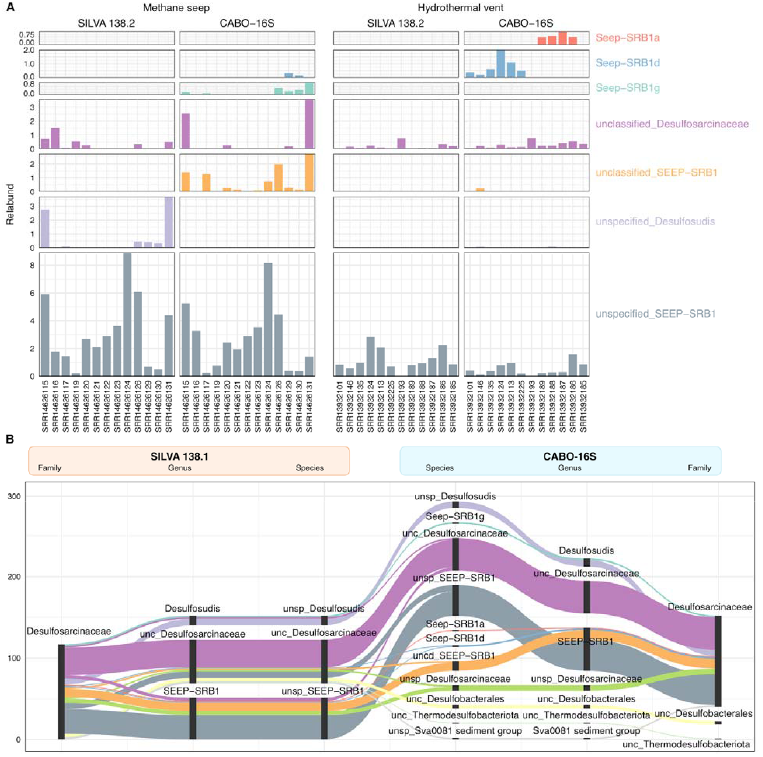
CABO-16S allowed classification of SEEP-SRB1 at higher resolution by adding in specific sequences. A) Relative abundance of SEEP-SRB1 and related taxa (colors, also subplot rows) at the lowest classified level. For the methane seep and hydrothermal vent datasets, the left subpanels show SILVA 138.2 classifications vs. CABO-16S classifications in the right subpanels. Y-axis is in % of each sample’s total number of reads. B) Alluvial plot showing classification of the same ASV sequences across databases. Each column is a different rank (Family through Species), with flows colored by the species assigned at the species rank with CABO-16S. Flow height reflects the number of ASVs. Abbreviations: unc, unclassified; unsp, unspecified; sed, sediment. Note that we distinguish between lack of annotation due to classification conflicts (unclassified) vs. due to incomplete annotation of reference taxa (unspecified), discussed above in context of Fig. 2.

### Challenges associated with adding custom sequences

While annotation for Seep-SRB1 subtypes could be achieved by the addition of known sequences with specific annotation, there remains a discrepancy between taxonomy, the hierarchical nomenclature system, and phylogeny, the evolutionary history of these clades. Other groups may require even more complicated approaches than the one we employed for SEEP-SRB1. The genera *Shigella* and *Escherichia* are emblematic of this conflict, as both are deeply convolved evolutionarily, yet the taxonomic legacy has continued to complicate reference database hierarchies [51, 52]. Other environmental groups like Synechococcus have similarly posed challenges to rectifying taxonomy and phylogeny for decades [53-55]. For such groups, adding sequences with a specific, phylogenetically correct hierarchy is unlikely to improve classification as the LCA method of resolving ambiguities assumes all sequences share the same hierarchy. Thus, all existing sequences would require similar reclassification following the desired phylogenetic framework and necessitate additional curation to ensure compatibility of the new taxonomic hierarchy with sequence similarity. Ultimately, the feasibility of rectifying phylogeny and taxonomy is limited by the signal embedded in the 16S rRNA gene, and while genome-based phylogenies and 16S rRNA phylogenies largely agree, they are not identical [56].

An additional barrier to improving resolution is errors or inconsistencies in taxonomic assignments, i.e. similar sequences with conflicting names; such errors are estimated to account for 1.5-17% of sequences in SILVA [57, 58]. Based on the assumption that the majority of sequences are correctly and consistently labeled, approaches like IDTAXA incorporate tools to identify and drop individual sequences that conflict with the majority of similarly-named sequences during training [32], and standalone tools also exist [58]. However, such approaches work best for taxa represented by many sequences, which is not always the case for environmental lineages in need of improved resolution. Classifier resolution and accuracy can also be improved by constraining the database to include only microbes specific to the habitat sampled as has been successfully done for many animal microbiomes [34, 59]. Such habitat-specific training sets are undoubtedly the best for focused research on a specific system. However, understanding the environmental context of specific taxa with broad distributions, like SEEP-SRB, necessitates approaches that maximize the resolution possible with general databases like SILVA.

## Conclusion

CABO-16S successfully combines bacterial and archeal 16S rRNA sequences from SILVA 138.2 and organellar 16S rRNA sequences from the PR2 database with customly selected sequences, resulting in increased taxonomic assignment of ASVs compared to SILVA 138.2. Specifically, with the addition of plastidal sequences from PR2, CABO-16S excels at identification of phototrophic eukaryotes in marine and lake waters without additional sequencing of both 16S and 18S primer sets. Although some 16S sequences, such as mitochondria from plants and algae, are still minimal and may impact classification of specific environments, CABO-16S reduces the number of unassigned phototrophs such that the remaining abundant sequence can be rapidly quarried. CABO-16S is also constructed such that custom sequences can be added. With addition of sequences from the polyphyletic clade of SEEP-SRB1, we saw increased taxonomic differentiation in hydrothermal vent sediment samples. This could help determine the likelihood of syntrophy in particular environments and increase our understanding of the communities which contribute to AOM. Although the addition of custom sequences must be done cautiously, considering the number of unspecified sequences within SILVA and the difficulty of constraining polyphyletic clades to the current taxonomy structure, this function of CABO-16S gives users the freedom to customize 16S taxonomic classification and potentially increase the understanding of specific environments. Finally, CABO-16S provides a framework that can be easily updated with the release of future versions of the SILVA and PR2 databases.

## Declarations

### Availability of data and materials

R scripts and workspace are available at https://github.com/emelissa3/CABO-16S. Custom sequences and other things are hosted permanently on Figshare (https://doi.org/10.6084/m9.figshare.27288090).

## Acknowledgements

The authors would like to acknowledge Dmitri Bilyk for logo design. We thank all the members of the Orphan lab for useful discussions. We also would like to thank the SILVA and PR2 teams for their work creating these databases.

## Authors’ contributions

EE and DU conceived the project and designed the implementation framework. EE and DU wrote R scripts and produced figures. SC helped with ARB. SC and RM helped obtain custom sequences. VJO provided computational analysis capabilities. EE and DU wrote the manuscript with input and final approval from all authors.

## Ethics approval and consent to participate

Not applicable

## Consent for publication

Not applicable

